# What does AlphaFold3 learn about antigen and nanobody docking, and what remains unsolved?

**DOI:** 10.1101/2024.09.21.614257

**Authors:** Fatima N. Hitawala, Jeffrey J. Gray

**Author notes:** For correspondence (FMS).

## Abstract

Antibody therapeutic development is a major focus in healthcare. To accelerate drug development, significant efforts have been directed towards the *in silico* design and screening of antibodies for which high modeling accuracy is necessary. To probe AlphaFold3’s (AF3) capabilities and limitations, we tested AF3’s ability to capture the fine details and interplay between antibody structure prediction and antigen docking accuracy. With one seed, AF3 achieves an 11.0% and 11.4% high-accuracy docking success rate for antibodies and nanobodies, respectively, and a median unbound CDR H3 RMSD accuracy of 2.73 Å and 2.30 Å. CDR H3 accuracy boosts complex prediction accuracy, with antigen context improving CDR H3 accuracy, particularly for loops longer than 15 residues. Combining I-pLDDT with Δ*G*_*B*_ improves discriminative power for correctly docked complexes. However, AF3’s 60% failure rate for antibody and nanobody docking (with single seed sampling) demonstrates necessary refinement to improve antibody design endeavors.

## Introduction

Antibodies (Abs) play a critical role in the immune system, and the development of antibody and nanobody therapeutics is a major interest due to their ability to target cancer, autoimmune, cardiovascular, and infectious diseases. Therapeutic advantages include their soluble nature, tunable affinity, high tolerance by the human body, manufacturability, and regulation history (Chungyoun and Gray, 2023). The antigen (Ag) binding interface of an antibody (nanobody) is composed of six (three) hypervariable loops, called the complementarity determining region (CDR) loops. The third loop on the heavy chain of the antibody (CDR H3) is particularly diverse and typically has the highest number of contacts with the epitope (Zhao et al., 2011). CDR loops sometimes undergo conformational changes upon binding to an antigen (Chu’nan Liu et al., 2024). Designing antibodies is challenging, primarily due to potential off-target effects (Chames et al., 2009) and the substantial time and resources required for developability testing (Chungyoun and Gray, 2023). Due to the flexibility and importance of antibody CDR loops, modeling structural movement and docking is highly valuable, and significant effort has been put into developing antibody and antibody-antigen complex structure predictors (Weitzner et al., 2017; Hummer et al., 2022).

Traditional Rosetta-based antibody-antigen docking algorithms use ensembles of homology models of antibodies and sampling of rigid backbones, loop conformation, and V_*H*_-V_*L*_ relative orientations (Weitzner et al., 2017). This general protocol has a 20% success rate for antibody-antigen docking, defined by a DockQ score > 0.23, which corresponds to an “acceptable” rating in CAPRI (Ambrosetti et al., 2020). Several other physics- and structure-based Ab docking methods have been published with similar performance (Sircar and Gray, 2010; Jiménez-García et al., 2018; Chen and Weng, 2002; Kozakov et al., 2017). While these methods are generalizable, the calculations are time consuming. Success rates are also limited by the ability to accurately model CDR H3.

Machine learning methods are faster and have significantly improved antibody-antigen and nanobody-antigen complex prediction. There are a variety of methods, with some focusing on only structure prediction or docking (Ruffolo et al., 2023; Abanades et al., 2023; Corso et al., 2023b; Jing et al., 2024; Chu et al., 2024), while others combine the tasks (Jumper et al., 2021; Verma et al., 2023; Jin et al., 2022). Tested architectures have included convolutional neural networks, transformers, diffusion models, and normalizing flow models for structure and complex prediction (Peng and Xu, 2011; Rao et al., 2021; Ruffolo et al., 2023; Corso et al., 2023b; Jin et al., 2022; Verma et al., 2023; Jing et al., 2024). While these methods have improved protein-protein docking success rates, AlphaFold2.3-Multimer (AF2.3-M) still had a poor 20% success rate for antibody-antigen docking (Ambrosetti et al., 2020; Harmalkar et al., 2024).

While AF2 and the the sub-series of models that were based off of the AF2 architecture (AF2.x-M) established a robust algorithm for predicting structure from processed sequence context, the limitations in docking and structure prediction of some protein families, ex. antibodies and nanobodies to antigens, have led to developments focused on improving the processed sequence context and sample diversity. Multiple Sequence Alignment (MSA) sub-sampling has been proven to extract conformational change information from sequence data (Rao et al., 2021), while massive sampling with increased diversity via tuned dropout rates performed the best in CASP15 (Wallner, 2023). AF3 is a culmination of many of these established methods.

Until AlphaFold3 (AF3), the highest reported success rate for antibody docking was 43% by AlphaRED (Harmalkar et al., 2024), a hybrid model using AlphaFold2-Multimer (AF2-M) predicted complexes and confidence measures with Rosetta-based replica exchange docking. Then in May 2024, with AF3 being trained on the same antibody dataset as AF2-M (Abramson et al., 2024; Jumper et al., 2021), DeepMind reported a notable 60% success rate for AF3 when 1,000 seeds were sampled.

To understand the source of improvement and where AF3 still has limitations, here we thoroughly assess AF3’s ability to dock antibody-antigen and nanobody-antigen complexes and predict unbound antibody and nanobody structures. To discern the effects of the limited experimental structures provided in the PDB, we study the interplay of the CDR H3 loop and Ab-Ag (and Nb-Ag) docking using structures from a redundancy-filtered bespoke dataset after the 2021 AF3 training cutoff. Using the DockQ score, we conduct an uncertainty quantification analysis to infer the likelihood of a correctly docked structure at varying confidence metric thresholds. We also determine the best combination of confidence metrics for antibody and nanobody complexes, and the utility of structure relaxation for decoy filtration. For reproducibility and standardized benchmarking, we make the benchmark data and code available at https://github.com/NooriFatima/AF3_AbNb_Benchmark.

## Results

### AF3 outperforms previous state-of-the-art antibody docking methods

To compare AF3 to previous state-of-the-art models, we first curated a benchmark set of bound and unbound antibodies and nanobodies (details in *Methods*). For every target sequence we ran three seeds in AF3 to account for the variance generated by the diffusion model. The AF3 paper reports an overall success rate of 60% for Ab-Ag docking, however this is with using 1000 seeds (Abramson et al., 2024). On the server we were limited by the number of jobs per day from running a greater number of seeds, and so we chose to focus the analysis on the quality of the predictions.

For consistency in comparison to the AlphaRED evaluation set, we chose the top-ranked decoy in the first seed (protocol in *Methods*). As seen in ***Figure 1***, while AlphaRED improves the percentage of acceptably docked structures to 43% compared to AF2-M, AF3 improves the overall docking quality by increasing the number of high-accuracy structures. AF2-M and AlphaRED both have negligible success rates for high-accuracy docking (DockQ ≥ 0.80) of antibodies, while AF3 has a considerably high accuracy success rate of 11.0% and overall success (DockQ > 0.23) rate of 38.1% (Figure 1A). For nanobodies, AF3 achieves a 11.4% success rate for highly accurate complexes, with a lower overall success rate (31.4%) (structures in **Figure 1—figure Supplement 1**).

**Figure 1.**
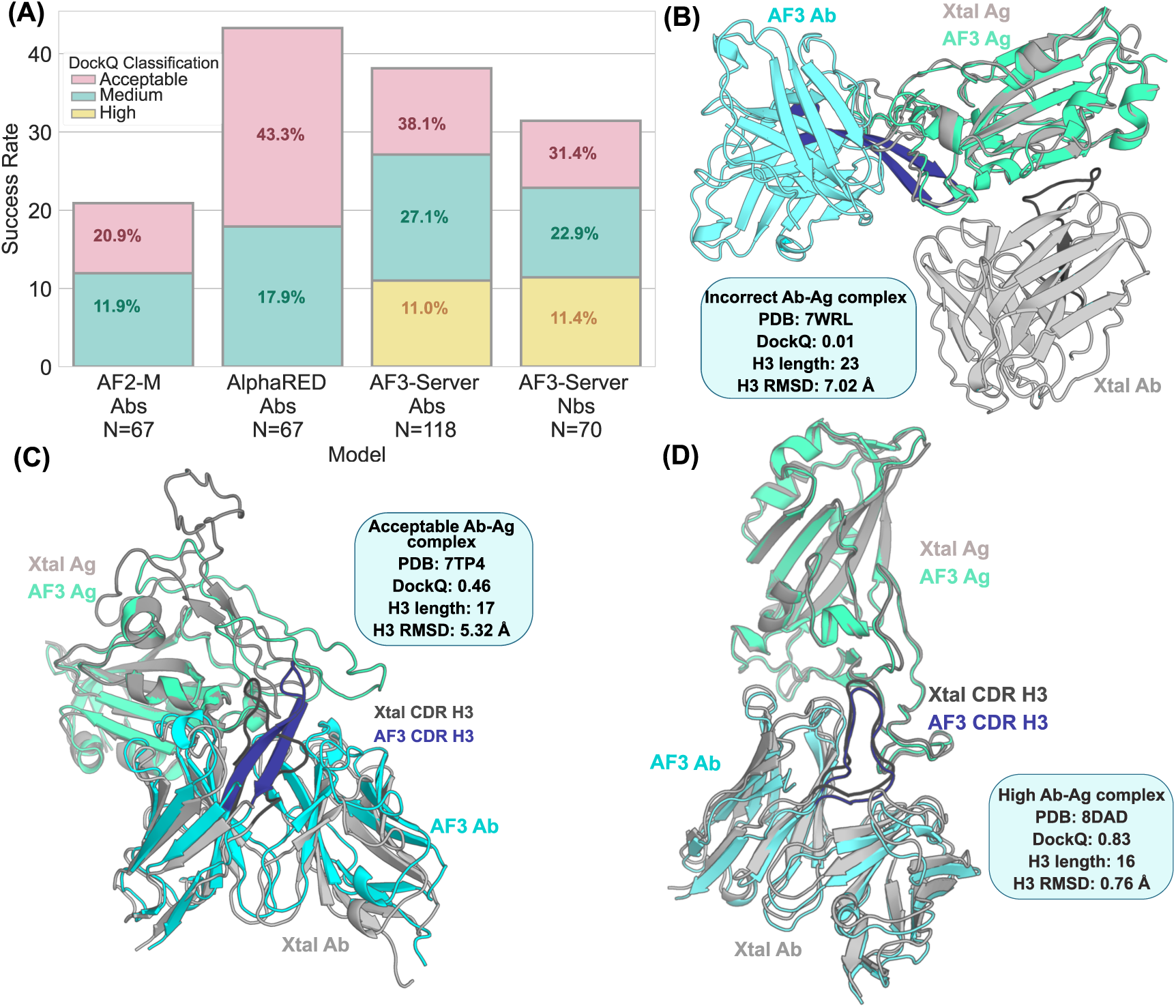
The success rates in antibody (Ab) and nanobody (Nb) docking for state-of-the-art models. (A) Performance of AF3 on antibody-antigen docking (N=118) and nanobody-antigen docking (N=70) with our curated dataset against AF2-M and AlphaRED (N=67) using AlphaRED’s evaluation sets (Harmalkar et al., 2024). DockQ scores for the bound antibodies and nanobodies are binned into incorrect, acceptable, medium, and high categories based on CAPRI classifications (Collins et al., 2024). (B, C, D) Ab protein complex structures of example incorrect, acceptable, and high accuracy predictions. Experimental crystal structures are gray, predicted antibody are blue, the predicted antigen is seagreen; crystal CDR H3 loop is dark grey and predicted CDR H3 loop is dark blue.

### Antibody structure prediction and antibody-antigen docking interdependently improve overall complex accuracy

The antibody maturation process improves binding affinity for an expressed antigen (Mishra and Mariuzza, 2018) so that a mature, unbound antibody can target the antigen by complementing the epitope (Conti et al., 2022). As the hypervariable H3 loop often makes the majority of contacts between the antibody and antigen (Zhao et al., 2011), correctly modeling the CDR H3 loop is pivotal in improving docking quality. This effect may be stronger for nanobodies, as they do not have a light chain with which to split the contacts to the antigen.

To understand the correlation between modeling the CDR H3 loop and docking accuracy, we measured the CDR H3 RMSDs and DockQ scores for AF3 predictions of antibody and nanobody complexes across three seeds and five diffusion decoys per target and visualized their joint distribution. In Figure 2, the predictions that are docked with high accuracy (DockQ ≥ 0.8, corresponding to the green box) are clustered within the bounds of CDR H3 RMSD ≤ 1.5 Å (the pink box). However predictions with CDR H3 RMSD ≤ 1.5 Å are scattered over the full range of possible DockQ scores. To describe this observation, we calculate the conditional probability, defined by Bayes’ Theorem: P(B|A) = P(A) and P(B)∕P(A). This probability quantifies the effect of one variable (either CDR H3 RMSD, or DockQ score) on the joint concentration of points defined by two variables (both CDR H3 RMSD and DockQ score).

**Figure 2.**
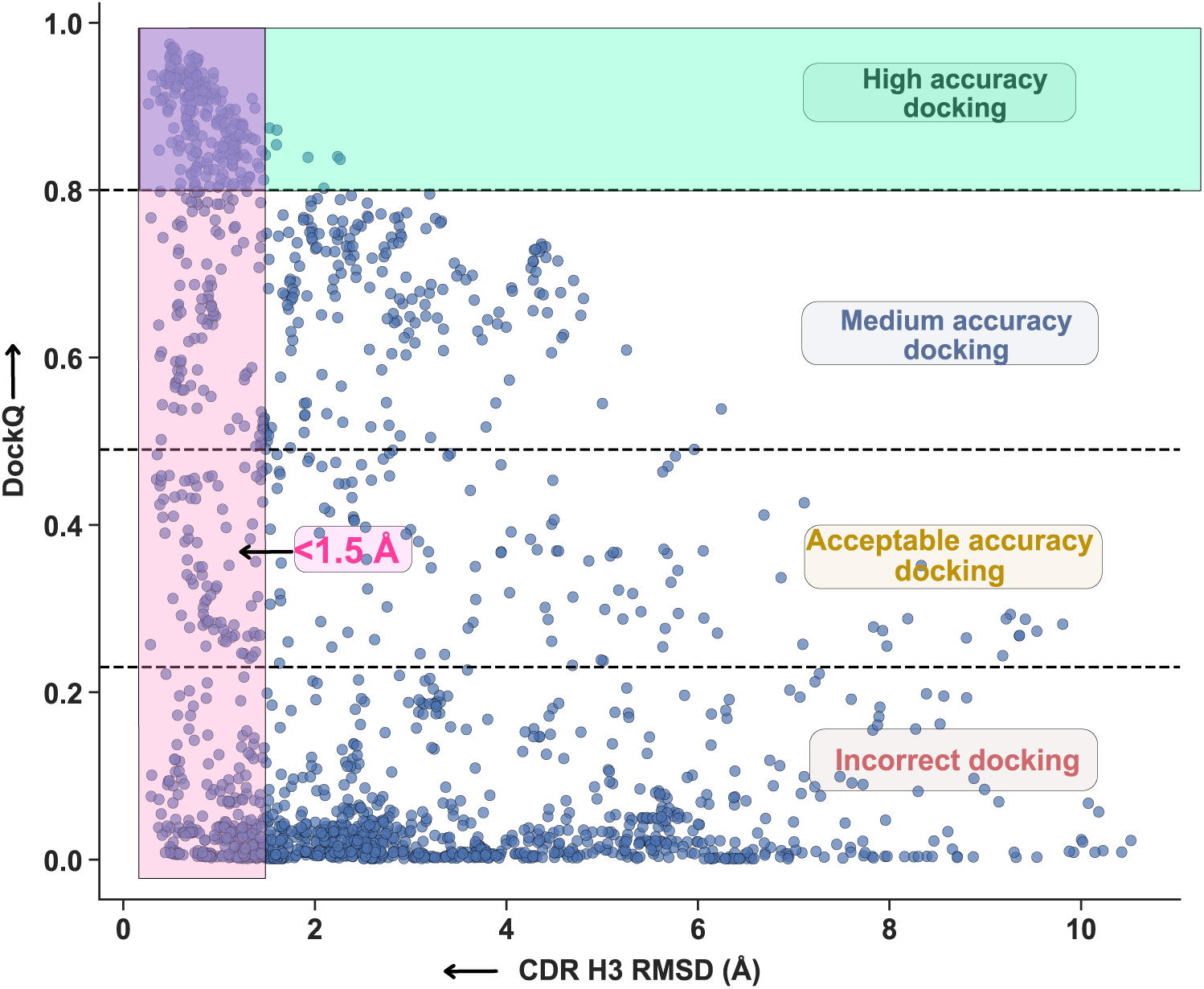
Distribution of DockQ scores versus CDR H3 loop RMSD of predicted antibody-antigen complexes. CAPRI classification zones marked, with the high-accuracy docking complex region shaded in green, the less than 1.5 Å CDR H3 loop RMSD region shaded in pink, and the intersection shaded in purple. The conditional probability of the CDR H3 loop RMSD being less than 1.5 Å given a highly accurate complex is the number of points in the intersection of both events (purple) over the number of total points with highly accurate docking (green). The conditional probability of a highly accurate complex given a less than 1.5 Å H3 loop RMSD is the number of points in the intersection (purple) over the number of points in the sub-angstrom H3 loop RMSD region (pink).

In this case, we can define the effect of CDR H3 RMSD on the joint concentration of points that have a CDR H3 RMSD ≤ 1.5 Å and a DockQ score > 0.8, and vice versa, the effect of DockQ score on the joint concentration of points that have a CDR H3 RMSD ≤ 1.5 Å and a DockQ score ≥ 0.8. Using the method in Figure 2 caption, we find that the p(DockQ ≥ 0.8 | CDR H3 RMSD ≤ 1.5 Å) is 27.8%, while the p(CDR H3 RMSD ≤ 1.5 Å | DockQ ≥ 0.8) becomes very high: 96.9%, implying that a correct CDR H3 loop is critical for high-quality docking predictions.

We then compared the probabilities of p(DockQ > X | CDR H3 RMSD ≤ 1.0 Å), where X is each DockQ score defining a class, against the p(CDR H3 RMSD ≤ 1.0 Å | DockQ > X) to find when p(DockQ > X | CDR H3 RMSD ≤ 1.0 Å) became larger than p(CDR H3 RMSD ≤ 1.0 Å | DockQ > X). This indicates when docking accuracy becomes a driving force for correctly predicting the CDR H3 loop structure. This change occurs relatively early, with 62% of sub-angstrom CDR H3 RMSD predictions having successfully docked complexes (DockQ > 0.23), compared to only 38% of successfully docked complexes having sub-angstrom CDR H3 RMSDs (Table 1, Table 2).

Visualizing the joint distribution of CDR H3 RMSD and DockQ scores for nanobodies (Figure 2—figure Supplement 1), the distribution of CDR H3 RMSD narrows towards being sub-angstrom RMSD as docking accuracy improves. In particular, we find that having sub-angstrom CDR H3 RMSD is heavily dependent on docking accuracy, as the percentage of sub-angstrom CDR H3 RMSD predictions improve from 13.4% to 64.7% as complexes go from incorrect to successful docking (Table 3,Table 4). Considering the importance of the CDR H3 loop for nanobodies, the stringency of CDR H3 accuracy for correct docking is justified. We forego this analysis for framework regions and remaining CDR loops for bound antibodies and nanobodies as their median RMSDs are subangstrom (Figure 2—figure Supplement 2).

### AF3 outperforms AF2.3-M, AF2-M, and IgFold in predicting unbound F_V_ structures

Considering the impact of CDR H3 loop accuracy on overall docking success, we sought to evaluate AF3’s predictive accuracy for unbound CDR H3 loops. That is, we examined the accuracy when predicting antibodies alone, without antigen. To compare AF3 against previous state-of-the-art structure prediction models, we use IgFold’s curated benchmark of 196 unbound antibody variable fragments and 70 unbound nanobodies (Ruffolo et al., 2023). For the comparison we used one seed with AF3 (seed = 1) and used the top-ranked antibody (ranking methodology in *Methods*) from the five decoys predicted.

Figure 3 shows CDR H3 accuracies. While AF2.3-M has the lowest median CDR H3 RMSD of the three previous state-of-the-art models at 2.73 Å, AF3 achieves 2.15 Å. IgFold and AF2.3-M perform similarly (2.87 Å for IgFold), with AF2-M performing worse (3.03 Å). Thus, AF3 has improved CDR H3 loop prediction by 0.58 Å (p≤0.05). The accuracy plateau reached by IgFold, AF2-M, and AF2.3-M led to questions about the limit of accuracy possible for the loop, considering its flexibility. AF3’s results prove that better predictions are possible. Further, a recent survey of 177 pairs of boundunbound antibody complexes reported that in 70.6% of antibody CDR H3 loops, binding-induced conformational changes are under 1 Å (Chu’nan Liu et al., 2024). Figure 3 also shows the CDR H3 RMSD between antigen-bound and unbound antibodies (Xtal B-U column); these measurements suggest that antibody loop prediction may still be able to be far more accurate.

**Figure 3.**
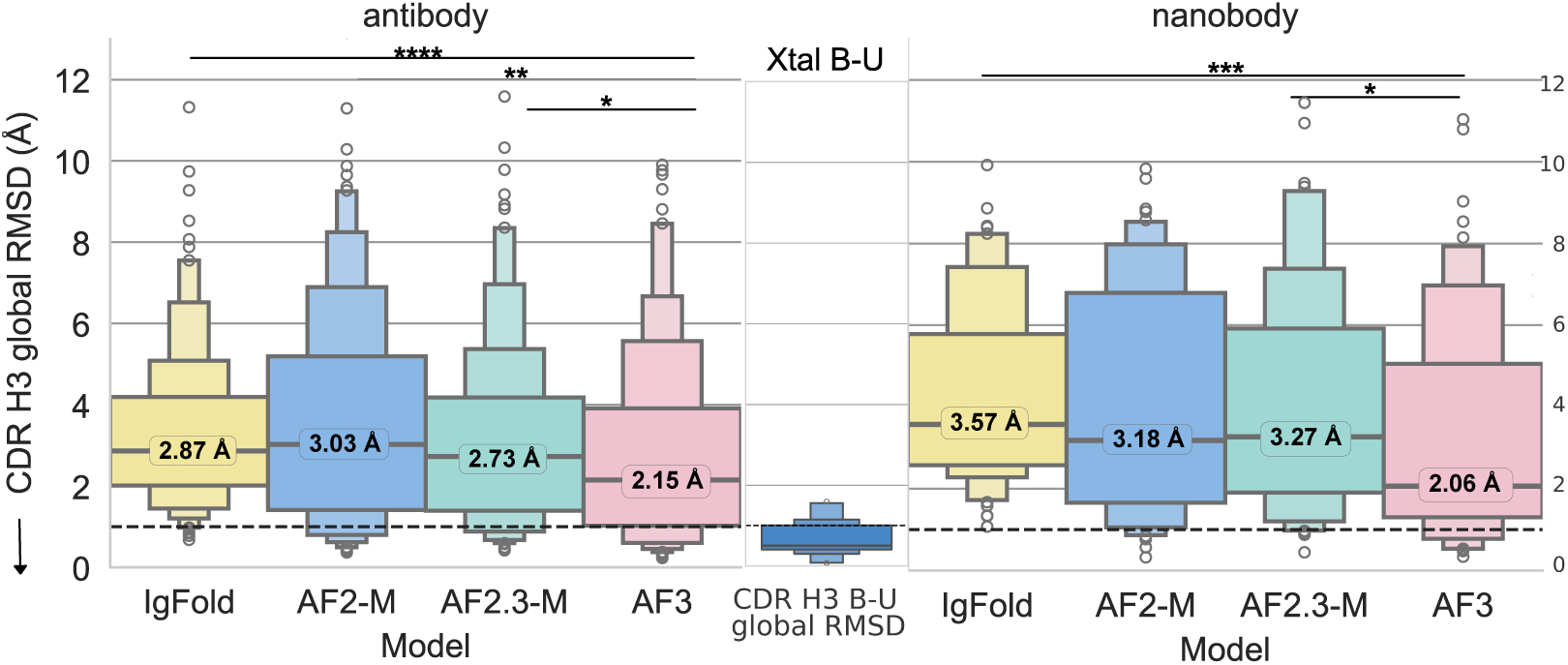
Performance of AF3 on predicting unbound CDR H3 loop structures of 196 antibodies and 70 nanobodies compared to previous models. The letter-value plot represents the traditional quartiles of the data, with additional boxes representing additional density. (A) AF3 improves upon the average median to 2.15 Å. For reference, the RMSD between unbound and antigen-bound antibody CDR H3 loops has a median of 0.5 Å. (B) While IgFold, AF2.0-M, and AF2.3-M have a median H3 RMSD for nanobodies above 3 Å, AF3 improves the median to 2.06 Å.

### Antigen context affects antibody CDR H3 loop prediction accuracy

To observe how the H3 loop’s structure is affected by antigen context, we compared the RMSD of all bound CDR H3 loops against all unbound CDR H3 loops, separately analyzing those of antibodies versus nanobodies. We divide the RMSD into the global H3 RMSD, calculated by superposing heavy chains, and the local H3 RMSD, calculated by superposing only the CDR H3 residues. The global RMSD captures the loop shape and placement, while the local RMSD represents the loop’s shape only. Instead of filtering for top-ranked decoys, we retained all five diffusion decoys to increase the power of the Mann-Whitney U test, since the effect of antigen context should remain consistent for all decoys, however we limit the analysis to a single seed (seed = 1) to avoid introducing noise from the variation seen between seeds. The medians of the CDR H3 loop RMSD vary by seed (Figure 2—figure Supplement 2), so we include the average median over the three seeds and their standard deviation in Table 5 and Table 6 for antibodies and nanobodies respectively.

Figure 4A shows that the global RMSD improves when given antigen context, from 2.73 Å to 2.05 Å (p=0.02), whereas AF3 predicts the local CDR H3 loops of bound and unbound Abs with the same accuracy (1.47 Å for unbound, 1.40 Å for bound). This implies that antigen context does not affect the predicted loop shape as drastically as it affects loop placement by providing spatial constraints. Previous studies report H3 loop RMSD increasing as loop length increases due to the degrees of freedom granted to the loop (Ruffolo et al., 2023). When antigen context is present to help constrain longer loops, as seen in Figure 4B, global AF3 CDR H3 accuracy improves significantly (p = 4.12×10^−^8) from 5.89 Å to 3.64 Å.

**Figure 4.**
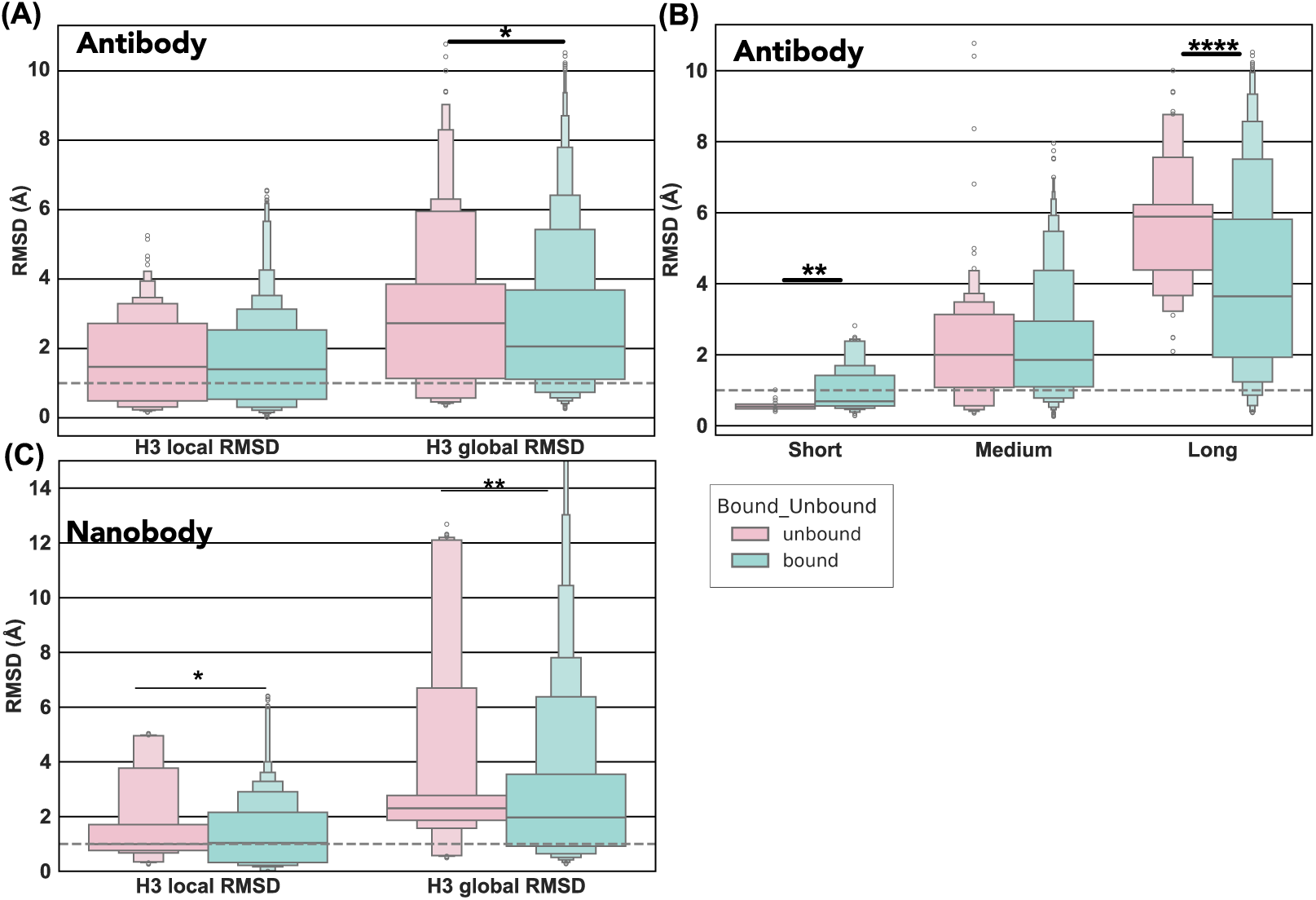
Effect of antigen context and loop length on CDR H3 loop prediction accuracy. (A) Effect of antigen context on antibody CDR H3 loop shape and position prediction accuracy. Bound structures (N = 1,770) predicted with antigen, unbound structures (N = 300) predicted with the antibody alone. Global RMSD calculated after superposition of the V_H_ domain; local RMSD calculated by superposing the loop residues only. (B) Effect of loop length and antigen context on CDR H3 loop prediction accuracy. Bound structures predicted with antigen, unbound structures predicted with the antibody alone. Short loops (N = 165) are defined as less than 10 residues, medium loops (N = 1,335) between 10 to 15 residues, and long loops (N = 570) longer than 15 residues. (C) Effect of antigen context on nanobody CDR H3 loop position and shape prediction accuracy. Bound structures (N = 1,005) predicted with antigen, unbound structures (N = 150) predicted with the nanobody alone. Global RMSD calculated after superposition of the V_H_ domain; local RMSD calculated by superposing the loop residues only. p ≤ 0.05 corresponds to *, p ≤ 0.01 corresponds to **, p ≤ 0.001 corresponds to ***, and p ≤ 0.0001 corresponds to ****

For nanobodies, we find global loop structure prediction improves (from 2.30 Å to 1.97 Å) given antigen context (p=0.008), as does local loop structure prediction slightly (from 1.04 Å to 0.99 Å) (p=0.05). The additional spatial constraints may affect nanobody CDR H3 loop structure more due to their larger average length compared to antibodies. We have insufficient data to make conclusions about the effects of loop length for nanobodies.

### A combination of predicted confidence interface metrics and Rosetta energies refines blind prediction scoring

Researchers need to know when to trust AF3 Ab-Ag and Nb-Ag structural models. Thus, we investigated confidence metrics and their respective cutoff values for scoring blind Ab-Ag and Nb-Ag docking. We use the top-ranked prediction for a single seed (seed = 1) for all targets to simulate the confidence filtration common in blind prediction ranking.

We first analyze ipTM-HA for both antibodies and nanobodies (outputted by AF3), because we have found that AF3 has a strong calibration of the ipTM (Figure 5—figure Supplement 1) as it is part of the training loss (Abramson et al., 2024). First, we generate distributions characterizing how likely a docked complex is to be correct (DockQ > 0.23) or incorrect (DockQ ≤ 0.23) depending on the confidence metric, which we calculate by binning the ipTM values through their total range, then generate counts of correct and incorrect docked complexes. As seen in Figure 5A-B, the likelihood of the complex being correct increases quickly when ipTM-HA ≥ 0.5 for antibodies, and ipTM-HA ≥ 0.4 for nanobodies (percentages listed in Table 7). Thus, we conclude that ipTM-HA ≥ 0.5 for antibodies, and ipTM-HA ≥ 0.4 for nanobodies are good cutoff values to discriminate correct docked complexes.

**Figure 5.**
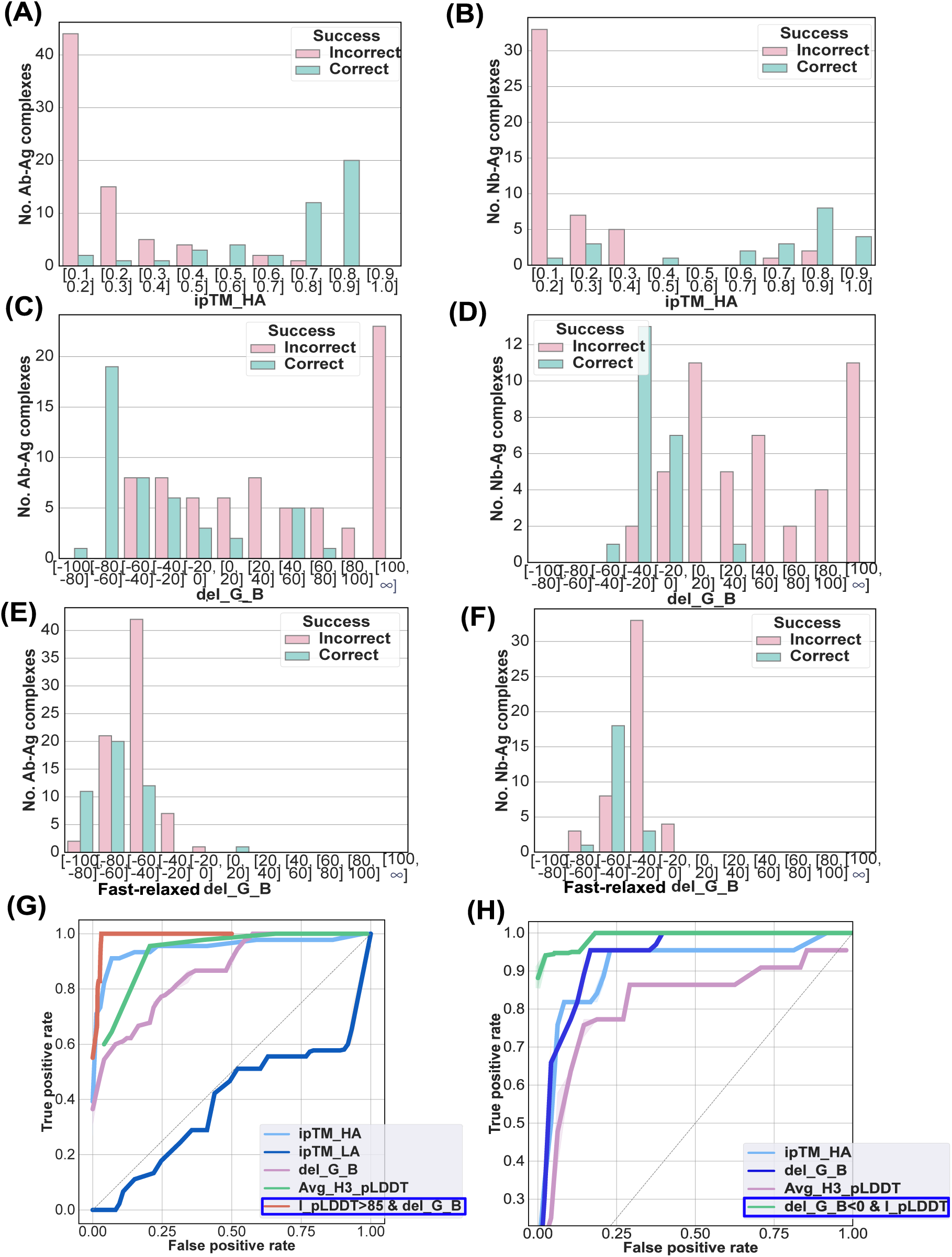
Optimized scoring metrics for Ab-Ag and Nb-Ag docking using AF3. (A) The number of Ab-Ag complexes that are correct (DockQ > 0.23) versus incorrect (DockQ ≤ 0.23) within a range of ipTM scores between the heavy chain and antigen. (B) The number of Nb-Ag complexes that are correct versus incorrect within a range of ipTM scores between the heavy chain and antigen. (C) The number of Ab-Ag complexes that are correct (DockQ > 0.23) versus incorrect (DockQ ≤ 0.23) within a range of Rosetta binding energy (ΔG_B_) scores. (D) the number of Nb-Ag complexes that are correct versus incorrect within a range of Rosetta binding energy (ΔG_B_) scores. (E) The number of Ab-Ag complexes that are correct versus incorrect within a range of Rosetta binding energy (ΔG_B_) scores after the complex has been FastRelaxed. (F) The number of Nb-Ag complexes that are correct (DockQ > 0.23) versus incorrect (DockQ ≤ 0.23) within a range of Rosetta binding energy (ΔG_B_) scores after the complex has been FastRelaxed. (G) AUROC curve comparing the _9 of 21_ performance of ipTM between the heavy chain and antigen (ipTM (H-A)) and between the light chain and antigen (L-A), versus the Interface predicted Local Distance Difference Test (I-pLDDT) for Ab-Ag complexes, ΔG_B_, and average CDR H3 pLDDT metrics. Also depicted is the best combined scoring metric, highlighted in

We conduct a similar analysis using Rosetta-based binding energies (ΔG_B_) (calculations in *Methods*), as it has been used to reliably rank protein structures (Alford et al., 2017). For antibodies, the likelihood of selecting a correct complex becomes greater than or equal to that of selecting an incorrect complex at ΔG_B_ ≤ −40 Rosetta Energy Units (REU), while for nanobodies, the cutoff is much higher (ΔG_B_ ≤ 0 REU) (percentages listed in Table 8). Due to the generative nature of the model, predictions from AF3 sometimes have steric clashes, (Abramson et al., 2024), so we considered the utility of Rosetta’s FastRelax in better filtering the output structures. After relaxing the complex (*Methods*), we recalculate the interface binding energy, and find that the range of energies has collapsed (Figure 5E-F). For antibodies, the new cutoff value (ΔG_B_ ≤ −60 REU) increases the false positive rate, making discrimination worse. Relaxing does not affect nanobodies, as at the optimal threshold (ΔG_B_ ≤ −40 REU) two complexes are false positives whether unrelaxed or relaxed.

As shown in Figure 5—figure Supplement 2A-B, we conduct the same analysis for the interface-predicted local distance test (I-pLDDT) as I-pLDDT is part of the training loss for the full model (Abramson et al., 2024), and it has been reported to be a stronger discriminator than ipTM for AlphaFold2-Multimer (Harmalkar et al., 2024; Yin and Pierce, 2024) (*Methods*). We find that the likelihood of selecting a correct complex is higher than that of an incorrect complex at I-pLDDT ≥ 85 for antibodies, and I-pLDDT ≥ 90 for nanobodies (percentages in Table 9).

We next analyzed the averaged pLDDT of the CDR H3 loop residues (avg. H3-pLDDT) (*Methods*), due to the interdependence we determined between CDR H3 accuracy and docking for antibodies and nanobodies. In Figure 5—figure Supplement 2C-D, we show that the optimal cutoff for antibodies is avg. H3-pLDDT ≥ 85, and avg. H3-pLDDT ≥ 90 for nanobodies (percentages in Table 10).

Combining scoring metrics provides the chance to tighten filtration and optimize the likelihood of sampling a correctly predicted complex. We simulate a two-metric filtration strategy by setting the first metric to the cutoff determined by the analysis mentioned above, then varying the threshold values of the second metric. To effectively compare the discriminative power of single and combined metrics, we used a normalized AUC (n-AUC) metric (*Methods*), with AUROC curves for single and the best combination metrics for antibodies and nanobodies highlighted in Figure 5G-H. To find the best discrimination thresholds, we find the point of the AUROC where the True Positive Rate (TPR) is the highest, and the False Positive Rate (FPR) is the smallest. For antibodies, first setting I-pLDDT ≥ 85, then again filtering with ΔG_B_ ≤ 65 REU (the combined metric has an n-AUC = 0.985) results in the best discrimination. Individually, each metric does not discriminate correct complexes as well (I-pLDDT n-AUC = 0.875, ΔG_B_ n-AUC = 0.861). Nanobodies are best filtered by first setting ΔG_B_ ≤ 0, then filtering again using I-pLDDT ≥ 73 (n-AUC = 0.992). Individual metrics again score lower (ΔG_B_ n-AUC = 0.937, I-pLDDT n-AUC = 0.889). Remaining metric combinations can be found in Figure 5—figure Supplement 2E-F.

## Discussion

AF3 builds on AF2’s performance by expanding capabilities towards general chemical structures through reducing emphasis on MSA processing and replacing the residue frame-based structure module with an atomic-precision diffusion model. We demonstrate that these architectural changes help AF3 dramatically improve the rate of high-quality docked antibody-antigen from 0 to 11.0%; however, AF3 (with one seed) still leaves 60% of the targets incorrect. The model has difficulty finding the correct antigen interface for a given antibody interface. The accuracy of the model has been reported to increase to 60% when evaluating 1,000 seeds (Abramson et al., 2024), a finding similar to the AF2 massive sampling study where artificial diversity was injected by tuning dropout rates (Wallner, 2023).

The impressive rate of high-accuracy docked complex predictions coincides with the interdependence we find between CDR H3 loop structure prediction quality and docking accuracy; AF3 demonstrates a 2.05 Å median global CDR H3 RMSD when given the antigen sequence (compared to 2.73 Å without antigen context). The docking-dependent improvement in CDR H3 loop prediction accuracy is likely due the multi-resolution property of the diffusion model, as both AF3 and DiffDock have found that larger noise scales used for training diffusion models correspond to the larger overall structure of the protein complex, while smaller noise scales correspond to local structure (Abramson et al., 2024; Corso et al., 2023a). As AF3 predicts structure and docks the complex simultaneously and has calibrated its confidence metrics to local and global scales through varying noise scales and the mini-rollout, both local and global structures are able to influence one another.

AF3 outputs a well-calibrated bespoke rank-score in the prediction’s confidence files, but often decoys will have the same rank score, causing difficulty in evaluation. We evaluated the robustness of the confidence metrics generated by AF3 against a benchmark of sequences and structures nonhomologous to the model’s training data to see if prediction ranking could be improved using additional metrics of CDR H3 loops and biophysical interface energies. We find that combining Rosetta-based binding energies with I-pLDDT for both Abs and Nbs improves discrimination.

When we started this study, we were limited by the number of jobs available per day on the AF3 server and the small dataset size due to data contamination concerns. Our results show that while AF3 is considerably more accurate in modeling antibody and nanobody structures and docked complexes than previous approaches, there remains room for improvement of the 60% failure rate for both antibody and nanobody docking for single seed predictions. The impressive CDR H3 loop structure prediction accuracy can also in principle still be improved relative to the limited overall ∼1 Å bound-to-unbound conformational changes in antibodies measured by Liu et al. (Chu’nan Liu et al., 2024). Researchers can now improve upon this baseline by running more seeds a local AF3 setup the authors of AF3 have made generously available (Callaway, 2024). Our curated benchmark set is available so that new methods replicating AF3 (Boitreaud et al., 2024; Wohlwend et al., 2024), and additional seed sampling in local AF3 can be tested. We now have more powerful tools than ever for Ab engineering, which open promising avenues to design and engineer Ab therapeutics and understand immunology better.

## Methods and Materials

### Individual structure evaluation set

To create an immunoglobulin structure prediction dataset, we pulled structures from SabDab (May 31, 2024 for Abs, June 4, 2024 for Nbs) (Dunbar et al., 2014), and temporally separated evaluation structures using the Sept. 30, 2021 training dataset date cutoff by AF3. We separated all antibody copies in the remaining PDBs, then conducted a quality filtration similar to (Chu’nan Liu et al., 2024), where we removed PDBs with a resolution ≥ 2.8 Å, or had missing residues in the CDR loops by comparing atomic sequence and sequence residues using the Bio.Seq python package (Cock et al., 2009). We used the Kabsch alignment algorithm to calculate the structural redundancy of variable heavy and light chains between pairs of structures (Ruffolo et al., 2023). We kept pairs of structures that had heavy and light chain RMSD > 1 Å, but only one representative out of a pair of redundant structures. Finally, we filtered the remaining structures based on sequence redundancy against each other and AF3’s training set with a sequence identity cutoff of 99% and 95% respectively. We conducted MSA alignment on the heavy and light chains separately via Abalign (Zong et al., 2023), and then a custom Python function to calculate the sequence identities. The number of structures at each step of this process is shown in Figure 6.

**Figure 6.**
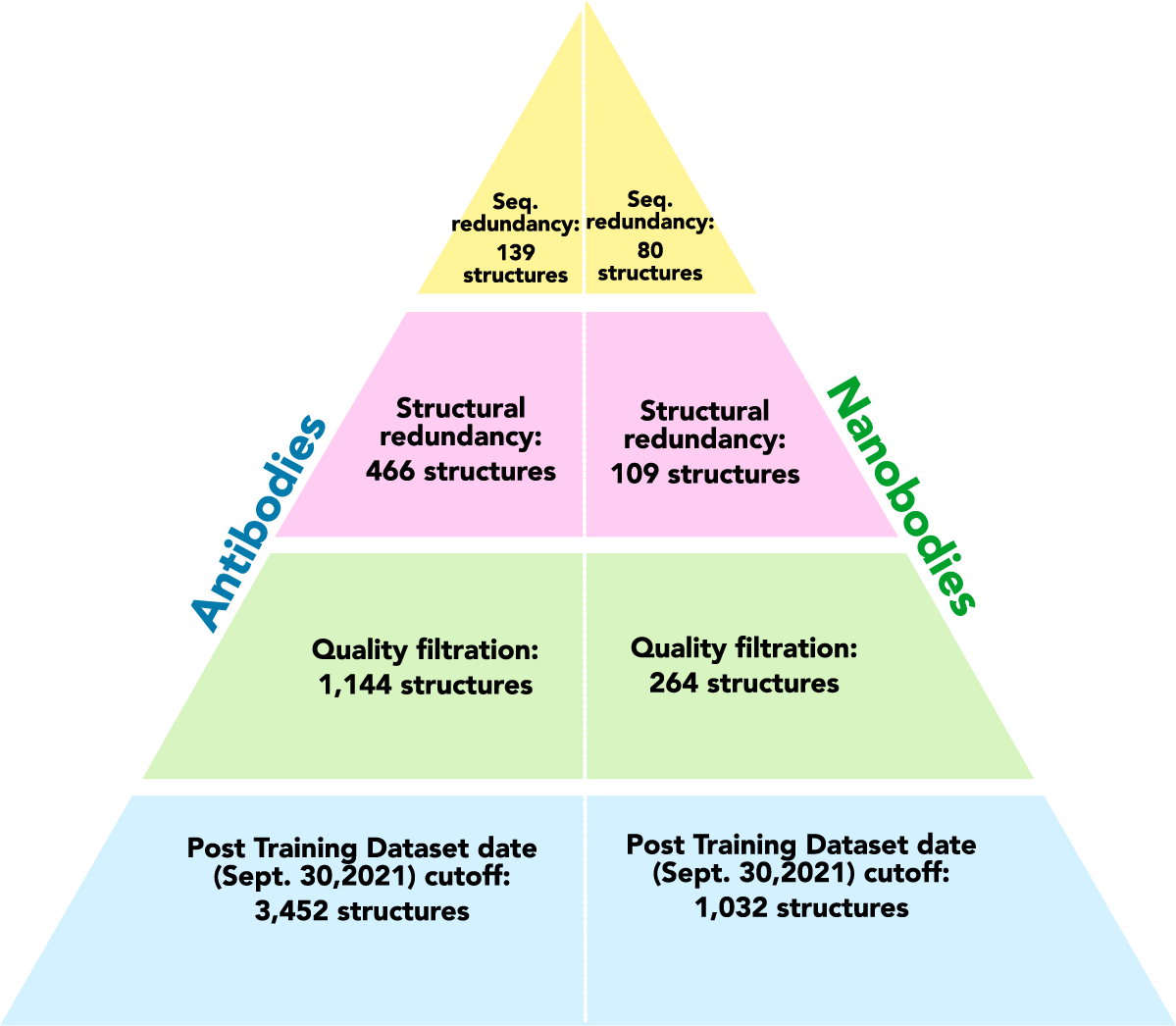
Dataset curation for evaluating antibody structure prediction and the number of structures at each step.

To prevent the RMSD calculations from being confounded by small hinge movements in the loop connecting the variable and constant regions, we cropped to the variable fragment region using a custom function. Finally, we renumbered the structures using the AbNum webserver (Abhinandan and Martin, 2008) using the Chothia scheme.

### AF2.3-M predictions

We used a local ColabFold installation with AlphaFold-Multimer version 2.3 downloaded from https://github. We predicted a single decoy for each target and did not use templates. Similar to the curated benchmark, we cropped the predictions to the variable fragment region using a custom function and renumbered the structures using the AbNum webserver (Abhinandan and Martin, 2008) using the Chothia scheme.

### AF3 predictions and ipTM extraction

We used the AF3 server (https://alphafoldserver.com) to generate decoys. The server generates five decoys (diffusion samples) per seed. To test the diversity produced by seeds, we predicted three seeds per target, with the seed number pre-set to either one, two, or three by using the JSON file upload option. We cropped the predictions to the variable fragment region using a custom function and renumbered the structures using the AbNum webserver (Abhinandan and Martin, 2008) using the Chothia scheme.

The TM-score measures topological similarity of two proteins, such that incorrect loop movements do not result in a severe scoring penalty, and is not dependent on protein size, making it more robust than the RMSD. The ipTM score is the predicted TM-score between two specified chains which compares the predicted complex to a theoretical complex (Zhang and Skolnick, 2004). AF3 outputs an ipTM score per chain. For our work we use either the score between the heavy chain and antigen, or light chain and antigen.

### Selecting the top-ranked decoy

We choose the top-ranked AF3 prediction per seed via the calculated rank score (a bespoke combination of ipTM, pTM, and disorder confidence measures) that AF3 provides in the confidence summary files with its predictions. If the rank score is equivalent for multiple predictions, we select the prediction with the highest ipTM-HA (as it is is applicable to both antibodies and nanobodies). If multiple predictions share a high rank score and ipTM, we select one from the pool randomly, as we aim to simulate blind prediction scoring.

### RMSD calculations

To calculate RMSD, we used PyRosetta with its AntibodyInfo class (Chaudhury et al., 2010). For calculating the CDR H3 global RMSD, we used the CDR backbone function, and for CDR H3 local RMSD, we extracted sub-poses for the CDR H3 loops also using the AntibodyInfo class, then calculated the Cα loop RMSD using the Rosetta Scoring class.

### Binding energy calculations

To calculate binding energies, we used PyRosetta’s InterfaceAnalyzer with the REF2015 score function (Chaudhury et al., 2010). We set the InterfaceAnalyzer to pack the complex before and after separating the binding partners to ensure that steric violations do not affect the rigid and flexible docking signals.

### Complex FastRelax

To remove steric clashes and improve predicted structure energies, we used the PyRosetta FastRelax method (Chaudhury et al., 2010) using the REF2015 score function, and a maximum of 100 iterations for speed.

### Figures and statistical analysis

We conducted Mann-Whitney-U and Pearson correlation statistical analyses using the Scipy package (Virtanen et al., 2020) in Python, and generated letter-value plots, scatter plots, and regression plots using the Seaborn package (Waskom, 2021) in Python. We used PyMol 3.0 (Schrodinger, Inc.) (Schrödinger, LLC, 2015) for protein structure visualization images.

### Area Under the Receiver Operating Characteristic (AUROC) curve

The AUROC curve is used to compare the ability of the confidence metrics to discriminate between correct (DockQ > 0.23) versus incorrectly docked (DockQ ≤ 0.23) complexes. We generate plot points by comparing the true positive rate at a particular threshold for the predictive metric to the false positive rate. As we vary the thresholds separating correct and incorrect cases for each metric, we generate a characteristic curve. The greater the area under the curve (AUC), the better the metric is at differentiating between the two classes. The AUC is calculated using the numpy trapezoidal function (Harris et al., 2020).

The true positive rates are calculated by dividing the number of true positive cases (DockQ > 0.23 and the metric > threshold), by the sum of true positives and false negatives (DockQ > 0.23 and the metric ≤ threshold). The false positive rates are calculated by dividing the number of false positives ((DockQ ≤ 0.23 and the metric > threshold)) by the sum of false positives and true negatives (DockQ ≤ 0.23 and the metric ≤ threshold).

Combined metrics are equivalent to a double filtering strategy. We use a union operator to combine an interface metric (ipTM or I-pLDDT) set at the optimized cutoff value with another measure, such as ΔG_B_ or avg. H3-pLDDT. For ipTM-HA, we use a cut-off value of 0.5 (antibodies), and 0.4 (nanobodies). For I-pLDDT, we use a cut-off value of 85 for antibodies and nanobodies despite the optimal threshold for nanobodies being 90 as there are no false positives above the cutoff, preventing AUROC generation. We also tested using ΔG_B_ as the first metric, then following with I-pLDDT to see if the order mattered. Many cases are pruned beforehand by the first metric, so combined metrics do not reach high FPRs, which means the maximum AUC they can reach will be < 1. Although the stringency of the combined metric is higher, the raw AUC will be lower. To combat this, we divide the raw AUC by the maximum FPR for that metric, resulting in the effective percent of discrimination.

### Interface and Average H3 pLDDT calculations

We calculate the I-pLDDT by averaging the residue pLDDTs for all residues within a 10 Å radius from the binding interface. We used the IMGT scheme in AbNumber (Dunbar and Deane, 2016) (IMGT scheme) with heavy chain sequences to extract pLDDTs of Cα residues for each H3 loop using the bfactor column through PyRosetta. We then averaged them in Python.

## Data and Code Availability

We make the curated benchmark dataset, PDB IDs used in prediction and their sequences, and the AF3 predictions available here: https://zenodo.org/records/14722282. The code we created for quick and standardized calculations and to visualize our results can be found here: https://github.com/Noori

## Acknowledgments

We thank the AlphaFold team for making the server available, Gray lab members for help in generating data from the server, and Samuel W. Canner and Emeline Haroldsen for comments on the manuscript. This work was supported by NIH R35 GM141881.

## Appendix

**Appendix 1—table 1.** Conditional probability of Ab docking accuracies given varying CDR H3 loop accuracies [p(DockQ > x | CDR H3 RMSD ≤ y Å)]

| DockQ Class \ H3 RMSD | <1 Å | <1.5 Å | <2 Å |
| --- | --- | --- | --- |
| High | 0.34 | 0.28 | 0.22 |
| High and Medium | 0.50 | 0.42 | 0.38 |
| High, Medium, and Acceptable | 0.62 | 0.53 | 0.48 |
| Incorrect | 0.26 | 0.35 | 0.40 |

**Appendix 1—table 2.**
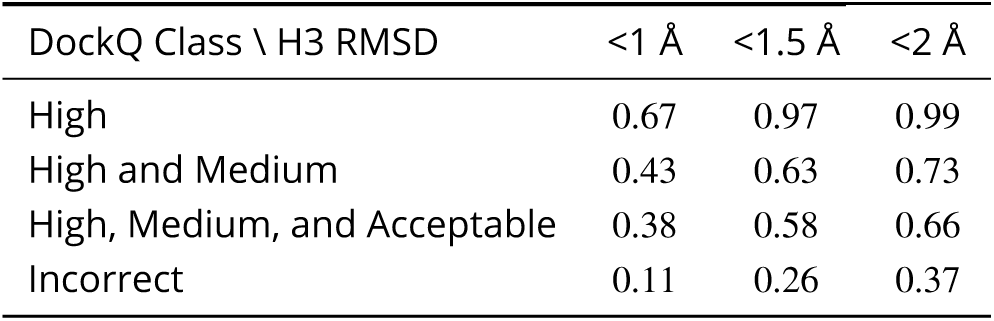
Conditional probability of Ab CDR H3 loop accuracies given varying docking accuracies [p(CDR H3 RMSD ≤ y Å | DockQ > x)]

| DockQ Class \ H3 RMSD | <1 Å | <1.5 Å | <2 Å |
| --- | --- | --- | --- |
| High | 0.67 | 0.97 | 0.99 |
| High and Medium | 0.43 | 0.63 | 0.73 |
| High, Medium, and Acceptable | 0.38 | 0.58 | 0.66 |
| Incorrect | 0.11 | 0.26 | 0.37 |

**Appendix 1—table 3.** Conditional probability of Nb docking accuracies given varying CDR H3 loop accuracies [p(DockQ > x | CDR H3 RMSD ≤ y Å)]

| DockQ Class \ H3 RMSD | <1 Å | <1.5 Å | <2 Å |
| --- | --- | --- | --- |
| High | 0.33 | 0.28 | 0.21 |
| High and Medium | 0.55 | 0.45 | 0.35 |
| High, Medium, and Acceptable | 0.64 | 0.59 | 0.48 |
| Incorrect | 0.32 | 0.37 | 0.43 |

**Appendix 1—table 4.** Conditional probability of Nb CDR H3 loop accuracies given varying docking accuracies [p(CDR H3 RMSD ≤ y Å | DockQ > x)]

| DockQ Class \ H3 RMSD | <1 Å | <1.5 Å | <2 Å |
| --- | --- | --- | --- |
| High | 0.85 | 0.99 | 1.0 |
| High and Medium | 0.77 | 0.90 | 0.93 |
| High, Medium, and Acceptable | 0.65 | 0.86 | 0.91 |
| Incorrect | 0.13 | 0.22 | 0.34 |

**Appendix 1—table 5.** Bound and Unbound top-ranked Ab CDR H3 loop RMSD (Å) variance by seed.

| Bound or Unbound \ average median and std. | median (Å) | std (Å) |
| --- | --- | --- |
| Unbound (N=21) | 2.80 | 0.23 |
| Bound (N=118) | 2.05 | 0.05 |

**Appendix 1—table 6.** Bound and Unbound top-ranked Nb CDR H3 loop RMSD (Å) variance by seed.

| Bound or Unbound \ average median and std. | median (Å) | std (Å) |
| --- | --- | --- |
| Unbound (N=10) | 2.28 | 0.12 |
| Bound (N=70) | 2.06 | 0.12 |

**Appendix 1—table 7.**
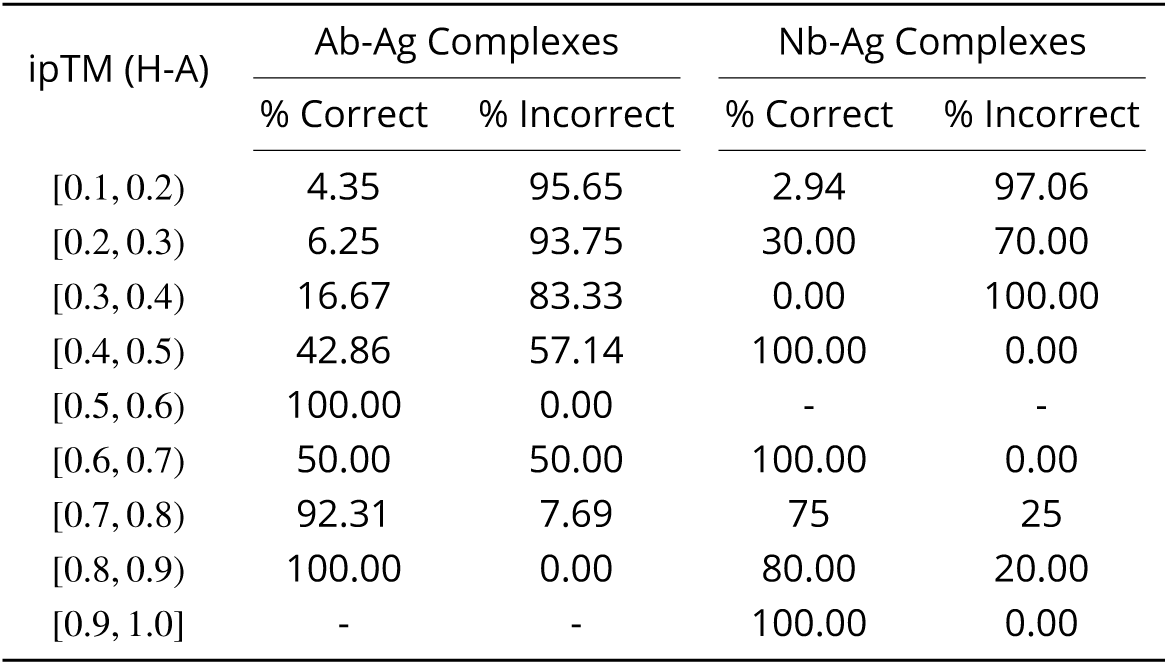
Percent of incorrectly versus correctly docked Ab-Ag and Nb-Ag complexes using ipTM between the heavy chain and antigen.

**Appendix 1—table 8.** Percent of incorrectly versus correctly docked Ab-Ag and Nb-Ag complexes using Rosetta binding energies.

| $\Delta G_B$ (REU) | Ab-Ag Complexes | | Nb-Ag Complexes | |
| --- | --- | --- | --- | --- |
|  | % Correct | % Incorrect | % Correct | % Incorrect |
| [-100, -80) | 100.00 | 0.00 | - | - |
| [-80, -60) | 100.00 | 0.00 | - | - |
| [-60, -80) | 50.00 | 50.00 | 100.00 | 0.00 |
| [-40, -80) | 42.86 | 57.14 | 86.67 | 13.33 |
| [-20, 0) | 33.33 | 66.67 | 58.33 | 41.67 |
| [0, 20) | 25.00 | 75.00 | 0.00 | 100.00 |
| [20, 40) | 0.00 | 100.00 | 16.67 | 83.33 |
| [40, 60) | 50.00 | 50.00 | 0.00 | 100.00 |
| [60, 80) | 16.67 | 83.33 | 0.00 | 100.00 |
| [80, 100] | 0.00 | 100.00 | 0.00 | 100.00 |
| [100, 1000] | 0.00 | 100.00 | 0.00 | 100.00 |

**Appendix 1—table 9.** Percent of incorrectly versus correctly docked Ab-Ag and Nb-Ag complexes using I-pLDDTs.

| I-pLDDT | Ab-Ag Complexes |  | Nb-Ag Complexes |  |
| --- | --- | --- | --- | --- |
|  | % Correct | % Incorrect | % Correct | % Incorrect |
| [70, 75) | 21.74 | 78.26 | 14.29 | 85.71 |
| [75, 80) | 13.64 | 86.36 | 7.69 | 92.31 |
| [80, 85) | 30.43 | 69.57 | 22.22 | 77.78 |
| [85, 90) | 100.00 | 0.00 | 33.33 | 66.67 |
| [90, 95) | 92.86 | 7.14 | 100.00 | 0.00 |
| [95, 100) | 100.00 | 0.00 | 100.00 | 0.00 |

**Appendix 1—table 10.** Percent of incorrectly versus correctly docked Ab-Ag and Nb-Ag complexes using averaged CDR H3 residue pLDDTs.

| H3-pLDDT | Ab-Ag Complexes |  | Nb-Ag Complexes |  |
| --- | --- | --- | --- | --- |
|  | % Correct | % Incorrect | % Correct | % Incorrect |
| [75, 80) | 3.70 | 96.30 | 20.00 | 80.00 |
| [80, 85) | 21.74 | 78.26 | 0.00 | 100.00 |
| [85, 90) | 61.11 | 38.89 | 42.86 | 57.14 |
| [90, 95) | 69.23 | 30.77 | 72.72 | 27.27 |
| [95, 100) | 100.00 | 0.00 | 75.00 | 25.00 |

---

**Figure 1—figure supplement 1.**
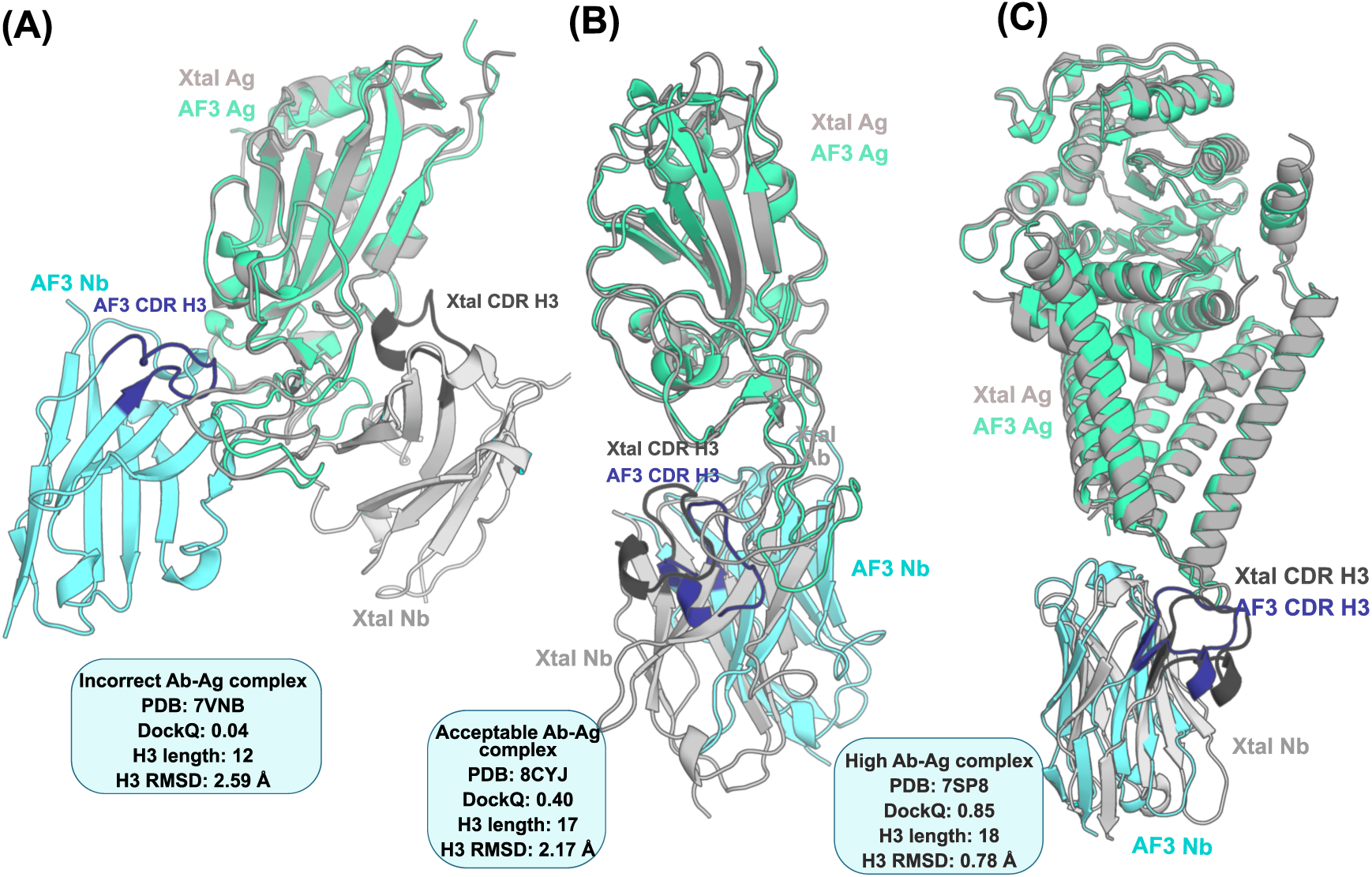
Examples of incorrect, acceptable, and high accuracy docked nanobody-antigen complexes. (A) incorrect nanobody-antigen complex compared to the crystal structure. The prediction samples the incorrect antigen interface and has a lower antigen structure prediction accuracy. (B) acceptably docked nanobody-antigen complex, where the antigen structure prediction accuracy is low, affecting the sampled paratope. The nanobody CDR H3 loop’s shape is correct, but the positioning is not. (C) The antigen and CDR H3 loop of the nanobody are correctly predicted, and the correct binding interface is sampled.

**Figure 2—figure supplement 1.**
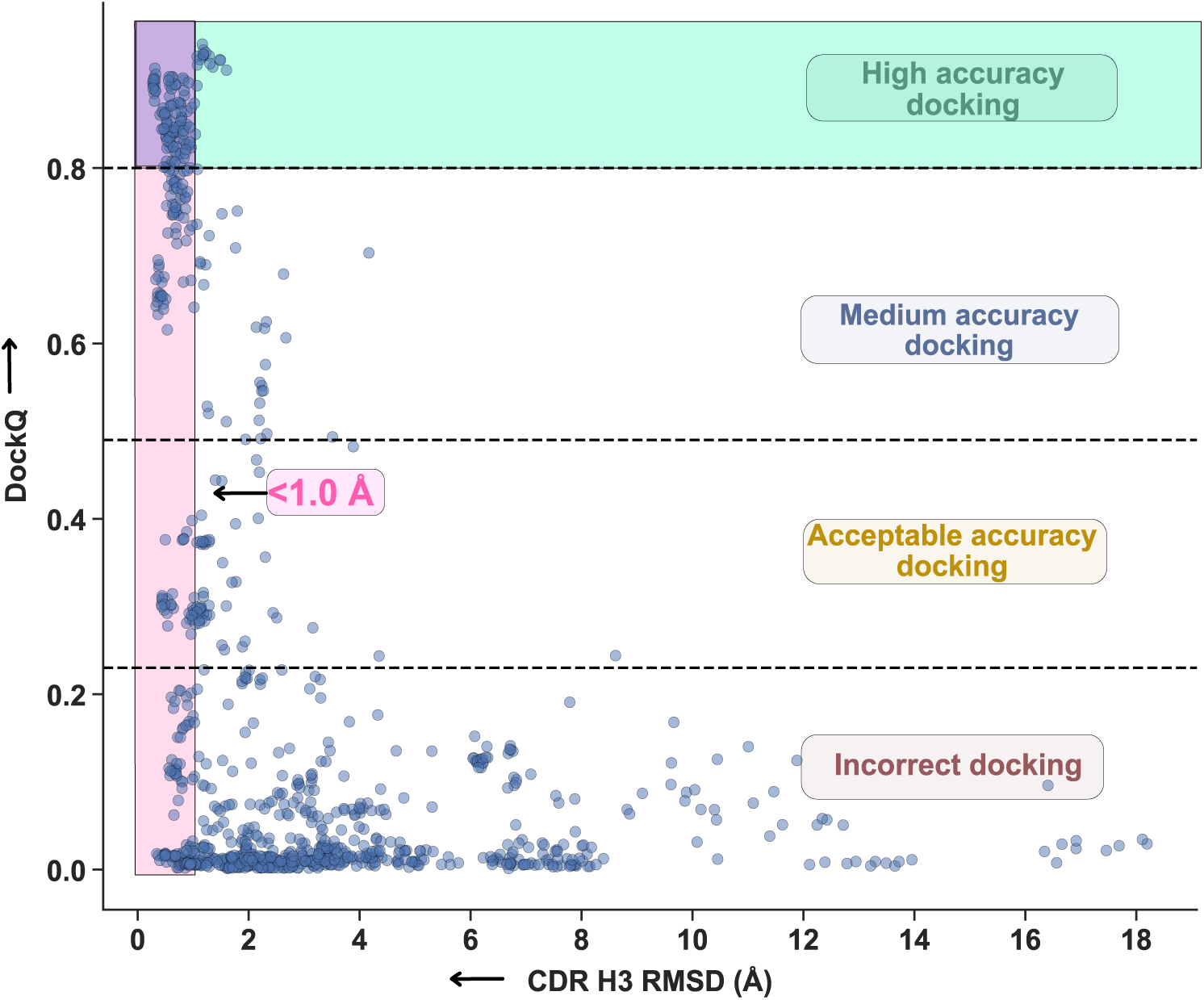
Distribution of DockQ scores versus CDR H3 loop RMSD for predicted nanobody-antigen complexes. CAPRI classification zones marked, with the high-accuracy docking complex region shaded in green, the sub-angstrom CDR H3 loop RMSD region shaded in pink, and the intersection shaded in purple. The conditional probability of the CDR H3 loop RMSD being sub-angstrom given a highly accurate complex is the number of points in the intersection of both events (purple) over the number of total points with highly accurate docking (green). The conditional probability of a highly accurate complex given a sub-angstrom CDR H3 loop RMSD is the number of points in the intersection (purple) over the number of points in the sub-angstrom CDR H3 loop RMSD region (pink).

**Figure 2—figure supplement 2.**
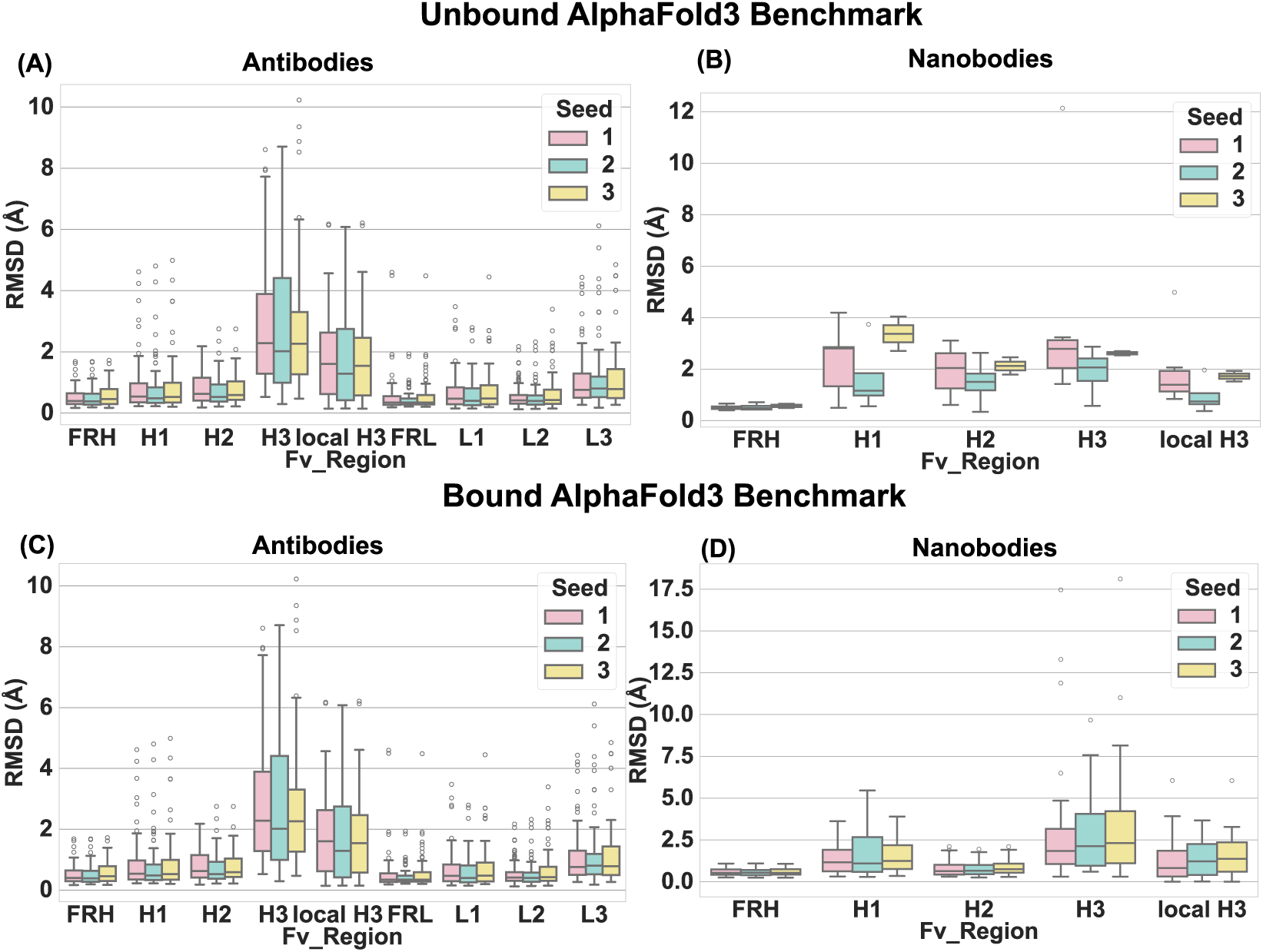
RMSD accuracies of framework regions and CDR loops for top-ranked bound and unbound antibodies and nanobodies per seed (A) unbound antibody RMSD accuracies. (B) unbound nanobody RMSD accuracies. (C) bound antibody RMSD accuracies. (D) bound nanobody RMSD accuracies.

**Figure 5—figure supplement 1.**
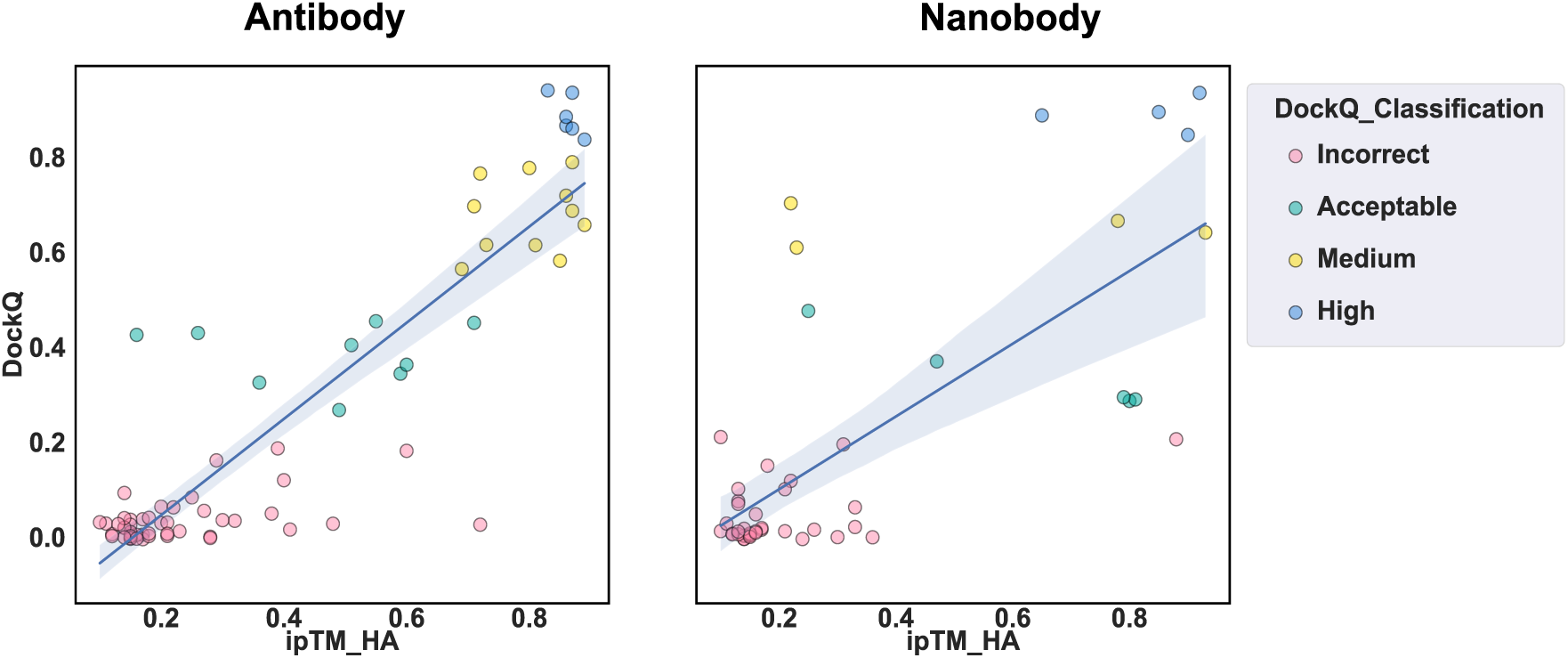
Correlation of ipTM between the heavy chain and antigen and DockQ score. (A) A linear correlation between the ipTM scores AF3 generates and the true DockQ score for top-ranked antibody-antigen complexes (R = 0.90, p = 5.60×10^−^27). (B) A linear correlation between the ipTM scores AF3 generates and the true DockQ score for top-ranked nanobody-antigen complexes (R = 0.73, p = 2.77×10^−^9).

**Figure 5—figure supplement 2.**
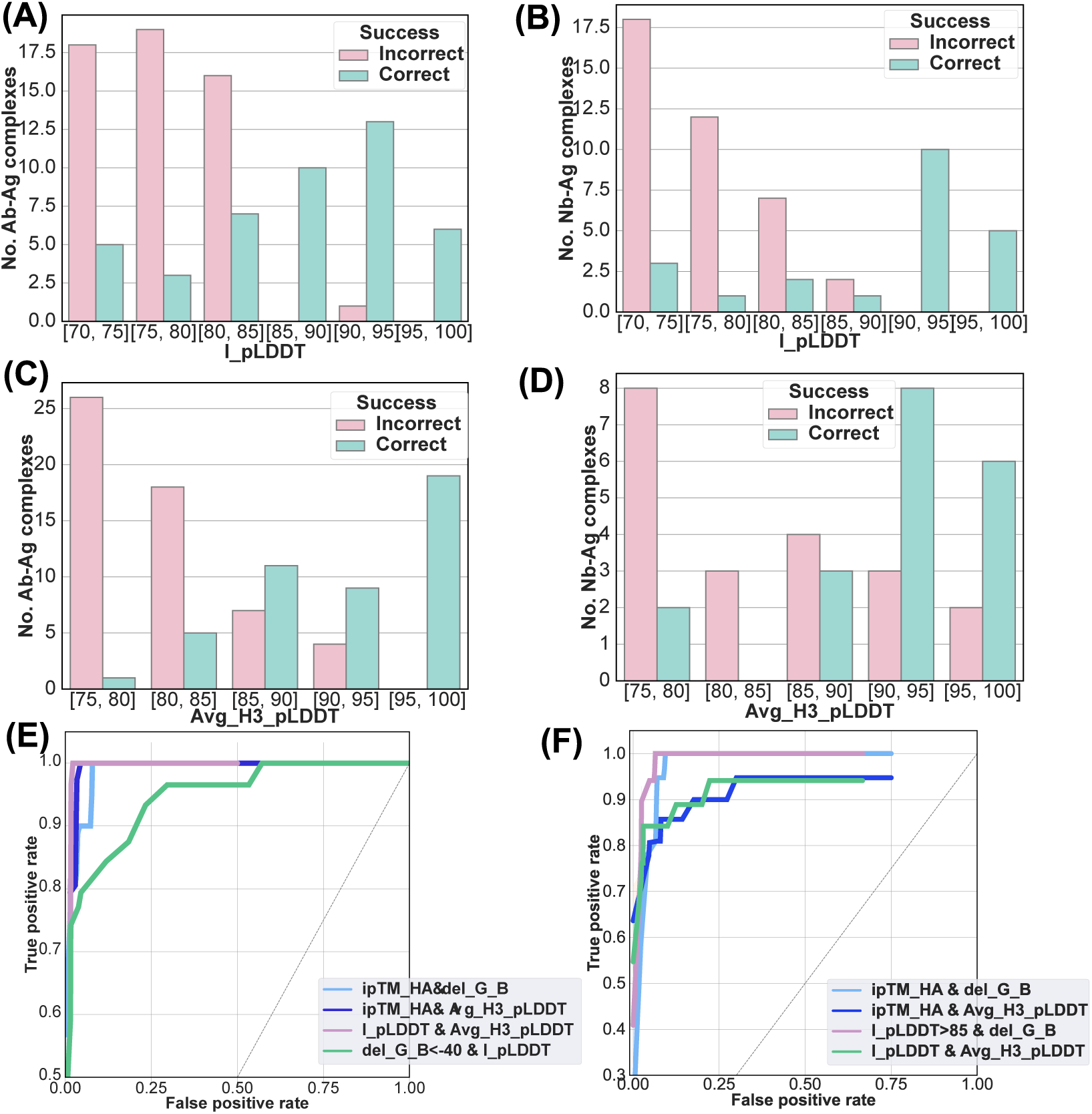
Quantifying ranking success probability of other widely used metrics for Ab-Ag and Nb-Ag docking. (A) The number of successful and unsuccessful Ab-Ag docks per range of I-pLDDT values and averaged CDR H3 loop pLDDTs. (B) The number of successful and unsuccessful Nb-Ag docks per range of I-pLDDT values and averaged CDR H3 loop pLDDTs. (C) AUROC curve of all other tested metric combinations for Ab-Ag complexes. (D) AUROC curve of all other tested metric combinations for Nb-Ag complexes.

## Notes

https://github.com/NooriFatima/AF3_AbNb_Benchmark

https://zenodo.org/records/14722282

### Competing Interest Statement

The authors have declared no competing interest.

### Summary of Updates

This revision summary has updated numbers for the previous analysis with greater stringency (DockQ, and antigen context sections), and extended analysis to include optimized protocols for scoring blindly predicted structures.

